# Mutations of schizophrenia risk gene *SETD1A* dysregulate synaptic function in human neurons

**DOI:** 10.1101/2024.10.20.619313

**Authors:** Xiao Su, Hanwen Zhang, Yan Hong, Qian Yang, Le Wang, Tiffany Le, Jiayi Liu, Lasya Cheruvu, Emily Labour, Siwei Zhang, Karla Mendez-Maldonado, Anat Kreimer, Hongjun Song, Guo-li Ming, Jubao Duan, Zhiping P. Pang

## Abstract

Schizophrenia (SCZ) is a complex neuropsychiatric disorder associated with both common risk variants of small effect sizes and rare risk variants of high penetrance. Rare protein truncating variants (PTVs) in *SETD1A* (SET Domain Containing 1A) show a strong association with SCZ; however, it remains largely unclear how rare PTVs in *SETD1A* contribute to the pathophysiology of SCZ. To understand the impact of *SETD1A* rare PTVs in human neurons, we CRISPR/Cas9-engineered five isogenic pairs of human induced pluripotent stem cells (iPSCs), with a recurrent heterozygous patient-specific PTV mutation c.4582-2delAG in two donor lines and a heterozygous frameshift mutation c.4596_4597insG (p. Leu1533fs) in three donor lines. These two mutations are predicted to cause a premature stop codon in exon 16 of *SETD1A*, leading to the loss of the conserved SET domain that is critical for its histone methyltransferase activity. We found that these presumably loss-of-function (LoF) mutations caused the *SETD1A* mRNAs to be degraded by nonsense-mediated decay (NMD), accompanied by a reduction of full-length SETD1A protein level in iPSCs. We then characterized the morphological, electrophysiological, and transcriptomic impacts of the *SETD1A^+/-^* LoF mutations in iPSC-derived human excitatory neurons induced by NGN2. We found that the *SETD1A^+/-^* exon-16 LoF mutations altered dendrite complexity, dysregulated synaptic transmission, and synaptic plasticity, likely by dysregulating genes involved in synaptic function. These results provide mechanistic insights into how *SETD1A^+/-^*exon-16 patient-specific LoF mutations affect neuron phenotypes that may be relevant to the pathophysiology of SCZ.

## Introduction

Schizophrenia (SCZ) is a chronic and severe mental disorder with an estimated heritability of 80%, affecting 1% of the population worldwide ^1^. Over the past decade, large-scale genome-wide association studies (GWAS) and exome sequencing have identified over 200 loci and around 120 risk genes associated with SCZ ^2–4^. However, the mechanisms by which genetic factors contribute to SCZ pathophysiology remain largely unclear, hindering the development of mechanism-based effective therapeutics.

Compared to common GWAS risk variants with small effect sizes, rare disease risk variants with relatively larger effect sizes and high penetrance provide important insights into disease biology. *SETD1A* (SET Domain Containing 1A) is one of the strongest risk genes for SCZ ^2,5–7^. Rare SCZ risk mutations, including protein-truncating variants (PTVs) in *SETD1A*, are also associated with intellectual disability, autism, and other neurodevelopmental disorders (NDD) ^8–12^. These rare disease-associated PTVs mainly include nonsense (i.e., protein codon stop-gain) mutations, splice variants, and protein-coding frame-shifting small insertion/deletions (Indels), presumably leading to protein loss-of-function (LoF). *SETD1A* encodes a component of the histone methyltransferase complex that plays essential roles in producing mono-, di, and trimethylation at histone 3 lysine 4 (H3K4) specifically ^13^. H3K4 trimethylation (H3K4me3) and H3K4me1 are canonic epigenomic marks of active gene transcriptional promoters and enhancers, respectively. Therefore, LoF in *SETD1A* may lead to an altered epigenetic landscape that causes gene expression dysregulation, contributing to the pathophysiology of SCZ. In support, histone methylation has also been suggested as one of the most enriched gene pathways in common variant-based GWAS of major psychiatric disorders ^14^, including SCZ ^15^. Moreover, most of the identified rare neurodevelopmental and SCZ-associated mutations of *SETD1A* appeared before the region encoding evolutionarily conserved SET domain (exon 17-19), which is critical for its histone methyltransferase activity ^7^. Nevertheless, it has also been reported that SETD1A has non-catalytic functions in the regulation of Cyclin-K and the DNA damage response ^16^, suggesting that SETD1A may function beyond an epigenetic regulator to mediate cellular function.

Haploinsufficiency in *SETD1A* has been studied in both animal and human stem cell models. Several mouse models of *Setd1a* deficiency using different strategies have been reported ^17–19^. These studies suggested that neuronal morphology, synaptic transmission, and gene transcriptions are affected by *Setd1a^+/-^* LoF. *Setd1a^+/-^*LoF targeting exon 4 in mice shows reduced axon branches and dendritic spines, increased neuronal excitability, and increased trend of excitatory synaptic transmission, affecting short-term synaptic plasticity ^17^. However, in another *Setd1a* LoF mouse model that targeted exon 7, a defective impaired excitatory synaptic neurotransmission was identified, which requires postsynaptic SETD1A to regulate the presynaptic glutamate release probability ^18^. Consistent with this, mouse neurons (both excitatory and inhibitory) carrying *Setd1a*^+/-^ LoF by targeting exon 15 also showed a defective synaptic release ^19^. Overall, these *Setd1a* haploinsufficiency mouse models suggested that synaptic transmission is likely one of the phenotypical manifestations of *Setd1a* LoF in neurons. Moreover, gene expression alteration is clearly affected by *Setd1a* LoF, likely due to altered H3K4me3 ^17–19^. In humans, individuals carrying *SETD1A^+/-^* LoF variants do not exhibit significant structural brain abnormalities or severe microcephaly ^7,20^, which indicates that impaired proliferation and differentiation of neural progenitors are unlikely to be the major underlying causes of the disorder. Similarly, it has been shown that *Drosophila melanogaster* SETD1A orthologue is essential in postmitotic neurons for memory formation of fly brain, indicating its role in post-development neuronal function ^20^. In addition to these animal models, *SETD1A*^+/-^ LoF is also studied in the human induced pluripotent stem cell (hiPSC)-derived neural model ^21,22^. In human-induced neuronal (iN) cells derived from one pair of isogenic hiPSCs carrying *SETD1A^+/-^* LoF mutation, it was found that reduced SETD1A expression levels lead to enhanced dendritic complexity and heightened bursting activity ^21^. These changes primarily manifest in glutamatergic neurons, as evidenced by altered gene sets linked to glutamatergic synaptic function in transcriptomic profiles reflecting these functional alterations ^21^. In human iPSC-derived cortical spheroid models, it was reported that reduced SETD1A levels lead to reduced neurite outgrowth, spontaneous activity, and altered metabolic capacity ^22^. Despite these studies, it appears that synaptic phenotypes in mouse models with *Setd1a^+/-^* LoF are inconsistent with very limited numbers of human iPSC lines carrying *SETD1A* mutations used in previous reports. Therefore, more extensive research is required to understand the synaptic and cellular phenotype in human neuronal models.

Of all the identified disease-associated *SETD1A* mutations ^17–19^, a two-base deletion at the exon 16 splice acceptor (c.4582-2delAG) is a recurring one that is most frequently carried by SCZ patients (four reported cases) and other NDD patients (three incidences) ^20^. The splice variant c.4582-2delAG is predicted to retain the intron 15, which leads to a short fragment of *de novo* protein sequence after exon 14 and a premature stop codon in exon 16 that causes loss of functional SET domain ^7^. To understand the impact of patient-specific mutation of *SETD1A* in a human context, we CRISPR/Cas9-engineered five isogenic pairs of iPSC lines carrying c.4582-2delAG or a frameshift mutation c.4596_4597insG (p. Leu1533fs) that mimics the LoF of this patient-specific mutation. We generated excitatory human iNs by ectopic expression of Ngn2 ^23^ and comprehensively characterized the transcriptional, morphological, and functional impacts of the patient-specific mutation in Ngn2-iNs. Overall, our results provide compelling evidence that supports *SETD1A^+/-^*exon 16 LoF mutations leading to nonsense-mediated decay (NMD) of *SETD1A* mRNA, altered neuronal dendrite complexity, and dysregulated synaptic function through transcriptional modifications.

## Materials and Methods

### iPSCs

Four iPSC lines (CD0000007, CD0000015 and CD0000019, and C11C) were selected for CRSIPR/Cas9 editing. C11C was reprogrammed from male fibroblast cells. CD0000007 (male), CD0000015 (female), and CD0000019 (male) were reprogrammed by Rutgers University Cell and DNA Repository (RUCDR)-NIMH Stem Cell Center from cryopreserved lymphocytes (CPLs) using integration-free Sendai virus method. All donors are control of European ancestry without large SZ-risk CNVs from the Molecular Genetics of Schizophrenia (MGS) study ^24^. As part of previous studies ^25,26^, the MGS donor-derived iPSC lines have been characterized for pluripotency by immunofluorescence (IF) staining of pluripotency markers and by RNA-seq-based Pluritest ^27^. The absence of chromosomal abnormality of these iPSC lines was confirmed by RNA-seq-based eSNP-Karyotyping ^25,26^, and all iPSCs were cultured at 37 °C, 5% CO_2_ on Matrigel® Matrix (Corning Life Sciences, CB-40234)-coated plates in mTeSR™ Plus medium (Stem Cell Technologies, # 100-0276), and passaged every 3-4 days using Accutase (Stem Cell Technologies, # 07920) The detailed procedure can be found in previous studies^28^.

### CRISPR editing of *SETD1A*

gRNA and ssODN for *SETD1A* editing were designed and gRNA was cloned into pSpCas9(BB)- 2A-Puro (PX459) V2.0 vector (Addgene, # 62988) following a previously published protocol (Ran et al., 2013). For editing, 24hr before transfection, 60-70% confluent iPSC culture was dissociated using Accutase (Stem Cell Technologies, # 07920) and replated onto matrigel-coated 60mm dish at 4.5 × 105 per well in mTeSR™ Plus with 5µM ROCK inhibitor (R&D systems, # 1254/1). The next day, DNA was transfected using LipofectamineSTEM (Thermofisher, # STEM00001) at 1:2 DNA:reagent ratio following vendor’s instructions in antibiotics-free mTeSR™ Plus with 5µM ROCK inhibitor. For the exon-16 1bp insertion editing, 4µg/well of Cas9 plasmid carrying the gRNA was introduced; for patient-specific 2bp deletion editing, 3µg/well of Cas9 plasmid carrying the gRNA and 3µg ssODN were used. 24hr post transfection, cells were selected in mTeSR™ Plus with 0.5 µg/ml puromycin. The media was refreshed with 0.25 µg/ml puromycin 48 hr post-transfection. 72hr post-transfection, the selection was withdrawn, and cells would be maintained in regular mTeSR™ Plus media for 7-10 days until colonies appeared and were large enough for genotyping. Colonies were manually picked onto 96-well plate and the genotypes at the mutagenesis site were confirmed by Sanger sequencing. All edited clones were confirmed by Sanger sequencing to be free of off-target editing at five top-ranking predicted off-target sites. The gRNA, ssODN, and primer sequences are available in supplementary table.

### Lentivirus production

Lentiviruses were produced as described ^29^ in HEK 293FT cells (ATCC) through co-transfection with 3rd generation helper lentivirus plasmid (pMDLg/pRRE, VsVG and pRSV-REV) using calcium phosphate transfections. Lentiviral particles were collected in mTeSR™ Plus medium, aliquoted, and stored at −80 °C. The following lentivirus constructs were used: (1) FUW-TetO-Ngn2-P2A-puromycin; (2) FUW-rtTA; (3) FSW-Venus; (4) pMax_GFP.

### Generation of iN Cells from Naive and Genetically Engineered Human iPSCs

Excitatory- (i.e. Ngn2-iNs) were generated as described ^23^. Briefly, iPSCs were dissociated by Accutase (Stem Cell Technologies, # 07920) and plated on six-well plate coated with Matrigel® Matrix (Corning Life Sciences, CB-40234) and mTeSR™ Plus medium containing RHO/ROCK pathway inhibitor Y-27632 (Stemgent, # 04-0012-02) or CEPT including Chroman 1 (MedChem Express, HY-5392), Emricasan (SelleckChem, S7775), Polyamine supplement (Sigma-Aldrich, P8483), and Trans-ISRIB (R&D Systems, 5284). At the same time of the plating, lentivirus cocktails prepared as described above (200 μl of Ngn2-contained virus and 200 ul of rtTA-contained virus per well of six-well plate) were added. On day 1, the culture medium was replaced with Neurobasal™ Medium (Gibco™, 21103049) containing doxycycline (MP Biomedicals, # 24390-14-5, 2 μg/ml), and doxycycline was maintained in the media for ∼10 days. On day 2 and 3, infected cells were selected by puromycin (Sigma-Aldrich, P8833, 1 μg/ml). On day 4, mouse glia cells were plated on Matrigel-coated coverslips (5 × 10^4^ cells/single well of 24 well plates), and the culture medium was replaced with Neurobasal™ Medium containing doxycycline (2 μg/ml) for recovery. On day 5, Ngn2-iNs (total of 2-3 × 10^5^ cells/well) were plated on a monolayer of mouse glial cells. On Day 6, media change occurred using Neurobasal medium with 5% FBS (fetal bovine serum, R&D Systems S11550) containing growth factors 10 ng/ml of BDNF (Peprotech, 450-02), GDNF (Peprotech, 450-10), NT3 (Peprotech, 450-03), B-27™ Supplement (Gibco™, 17504044), 1% GlutaMAX™ (Gibco™, 35050061), and Cytosine β-D-arabinofuranoside (AraC, Sigma-Aldrich, C1768, 2 μM), which was added to stop glial cell division. Morphological, biochemical, and functional analyses were conducted at 5 to 6-week-old. Each of the morphological and functional analyses was conducted from at least two or three independent culture batches for each isogenic pair.

### Immunohistochemistry

Primary antibodies were diluted in blocking buffer (1% goat serum (Gibco™, 16210072), 0.25% Triton X-100 and 4% BSA) at 4 °C and incubated overnight. Coverslips were washed 3 times with blocking buffer and then incubated with secondary antibody (1: 500) at RT for 1 hr. Following three washes with PBS, coverslips were mounted onto glass microscope slides with mounting medium containing DAPI (Sigma-Aldrich, F6057). The following antibodies were used: SOX2 (Millipore Sigma AB5603, 1:1000), Oct4 (Millipore Sigma MAB4401/4419, 1:1000), Tra-1-60 (Millipore Sigma MAB4360, 1:1000), MAP2 (Sigma-Aldrich M1406, 1:1000; Millipore Sigma AB5543,1:1000), Synapsin-1 (E028, 1:3,000; SYSY 106011,1:1000). At least two-three independent culture batches for each isogenic pair were used for all experiments.

### Sparse transfection in Ngn2-iN cells

To quantify changes in neurite outgrowth, Ngn2-iNs were sparsely transfected with FSW-Venus or pMax-GFP construct at day 40 using calcium phosphate-based transfection. Briefly, per one well of a 24-well format, 0.5 µg of GFP plasmid, 2.5M CaCl_2_, and molecular-grade water was mixed and added dropwise to 2X HBS (280 mM NaCl, 1.5 mM Na_2_HPO_4_, 50 mM HEPES) on a gentle vortex, and transfection mix incubated at room temperature for 10 min. After washing wells three times with MEM+++ (MEM containing 0.5 mM CaCl_2_, 1 mM MgCl_2_, and 4% sucrose to minimize coverslip peeling), transfection mix was added dropwise to each well and incubated for 30 min at 37°C. Wells were washed three times with MEM, and transfection efficiency was confirmed 48-72 hours later.

### Image acquisition and quantification of neurite outgrowth

All images were taken with a Zeiss LSM700 laser-scanning confocal microscope. Neurite outgrowth was captured using a 20x objective at 512 x 512-pixel resolution with tile scan and Z-stack function. Specifically, the filamenting modules of Imaris software (Bitplane version 9.5) were used to quantify changes in neurite outgrowth (metrics: number of branch points, total neurite length and *Sholl* analysis). Statistical analysis was performed using Student’s t test comparing control to each individual mutation (*p < 0.05, **p < 0.01, ***p < 0.001; nonsignificant comparisons were not indicated).

### Synaptic puncta density analysis

Immunofluorescence was performed on iNs at day 35-42 with dendrite marker MAP2 and synaptic marker Synapsin-1. Synapse formation was imaged using a Zeiss LSM700 laser-scanning confocal microscope with a 63x objective at 1024×1024 pixel resolution. All images were acquired using Z stack maximal projection in Zeiss Zen blue software. Correlated synaptic puncta number was quantified by Intellicount as reported previously ^30^. Primary processes were blindly counted and quantified.

### Western blot analysis

For iPSCs culture on 6-well plate, the cultures were washed twice with ice-cold PBS and then cells were lysed by 100 ul RIPA buffer (50 mM Tris-HCl pH7.5, 150 mM NaCl, 1% NP-40 (Igepal CA-630), 0.5% Sodium deoxycholate, 0.1% SDS, 1mM DTT, 1X Protease Inhibitor cocktail and phosphatase inhibitor) for each well. The protein concentration was quantified by BSA kit (Thermo scientific, #23227). For induced neuron culture on coverslips, at day 35-42, the cultures were washed once with ice-cold PBS. The proteins were directly collected by 2X SDS loading buffer (4% SDS, 125 mM Tris HCl (pH6.8), 20% glycerol, 5% 2-mercaptoethanol and 0.01% bromophenol blue) ^31^. Protein samples were then denatured by heating to 95 °C for 5 min. The denatured protein samples were allowed to cool down and equal amounts of samples were analyzed by 4–15% or 7.5% TGX gels (Bio-Rad #4561084DC, #4561023DC) for sequential western blots on the same membranes. The protein from the gel was transferred to a 0.45 μm or 0.22 μm nitrocellulose membrane (Bio-Rad #1620115) using the Bio-Rad transferring system. The nitrocellulose membrane was blocked in 10% nonfat milk for 1 h at room temperature. After the blocking step, membranes were incubated with primary antibody overnight at 4 °C. The following day, the blots were washed three times with Tris-buffered saline with 0.1% Tween 20 (TBST) before adding horseradish peroxidase-conjugated secondary antibody at room temperature for 1 h. Primary antibodies catalog number (N-SETD1A Invitrogen PA5-35975; SETD1A Abcam AB243881; H3K4me3 Cell Signaling 9751S; H3 Sigma-Aldrich H0164; VCP K330). Protein bands were visualized by the addition of Clarity Western ECL Substrate (Bio-Rad #1705060). The density of the bands was quantified using ImageJ software. Beta-actin was measured as a loading control.

### Electrophysiological Recordings

Whole-cell patch-clamp electrophysiology for Ngn2-iNs was performed as described ^28^. For miniature postsynaptic current and current-clamp recordings, K-gluconate internal solution was used, which consisted of (in mM): 126 K-gluconate, 4 KCl, 10 HEPES, 0.05 EGTA, 4 ATP-magnesium, 0.3 GTP-sodium, and 10 phosphocreatine. For miniature postsynaptic current recording, a Cs-based internal solution was used, which consisted of (in mM): 40 CsCl, 3.5 KCl, 10 HEPES, 0.05 EGTA, 90 K-gluconate, 1.8 NaCl, 1.7 MgCl_2_, 2 ATP-magnesium, 0.4 GTP-sodium, and 10 phosphocreatine. Miniature excitatory postsynaptic currents (mEPSCs) were observed for iN cells in voltage-clamp mode with a holding potential of 70 mV in the presence of 1 μM tetrodotoxin (TTX). For evoked recording, a Cs-based internal solution was used with QX314 (5 mM). The bath solution contained (in mM): 140 NaCl, 5 KCl, 10 HEPES, 2 CaCl_2_, 2 MgCl_2_, 10 Glucose, pH 7.4. To elicit action potentials, a series of increasing amounts of current was applied to the cell in 10-pA increments across 19 independent sweeps (−50 pA to +130 pA). Each step lasted for 1 s, and action potential kinetics were analyzed in ClampFit (11.1) for the first triggered AP observed at the lowest state of the current injection. All cell culture recordings were conducted in the whole-cell configuration at room temperature.

### RNA isolation and RNA sequencing

For qPCR reactions, cells from hiPSCs and iNs cultures were lysed in RLT plus buffer, and total RNA was purified with the RNeasy mini-Kit (QIAGEN). For RNA sequencing, the cultured iNs were lysed in RLT plus buffer (QIAGEN) and subjected to RNA sequencing (RNA-seq) at GENEWIZ from Azenta (South Plainfield, NJ) and Novogene (Sacramento, CA).

### Quantitative RT-PCR

Purified RNA was eluted in 35 uL per sample. Purified total RNA was reverse-transcribed and PCR-amplified using cDNA reverse transcription kit (Invitrogen #11755050). The qPCR reactions were conducted with PowerUp SYBR Green Master Mix for qPCR kit (Applied Biosystems, # A25742). The mRNA levels were quantified by real-time PCR assay using the Quantstudio 3 System and RQ analysis software (Thermo Fisher Scientific). Within each plate, the housekeeping gene GAPDH was included as an internal control.

### Gene Expression Analyses

#### Bulk RNA-sequencing

For samples with c.4596_4597insG mutation, RNA-seq files were provided in the format of 2 × 150 bp paired-end fastq files from GENEWIZ from Azenta. Briefly, cDNA libraries were prepared with the polyA selection method with Illumina HiSeq kit and customized adapters. 20–30M paired-reads were recovered from each sample at the rate of ∼350 M raw read-pairs per lane on an Illumina HiSeq 2000. No Trimmomatic-based adapter-trimming was performed, as the sequencing facility had performed pre-cleaning on raw reads. Raw files were subsequently mapped to human hg38 genome (GRCh38p7.13) with STAR v2.7.0 and GENCODE v35 annotation file.

For samples with c.4582-2delAG mutation, paired-end reads per sample were collected using NovaSeq PE150 platform (Novogene) with polyA enrichment. Quantified libraries were pooled and sequenced on Illumina NextSeq500 platforms by Novogene. Approximately 40-60 million paired reads were generated for each sample. Reads were aligned to the human hg38 reference (GRCh38p7.13) genome by HISAT2 (v.2.1.0) ^32^.

#### Differential gene expression analysis

DESeq2 (v1.28.1) was used to perform DEG analysis for raw count values of samples with the design formula as ‘‘design = ∼ cell_lines + genotypes’’, with ‘‘cell_lines’’ as the key of paired samples and ‘‘genotypes’’ as the key of genotype annotations. Significance test results from DESeq2, including log2 fold changes and FDR adjusted p values, were used for downstream analysis. Ensembl ids were transferred to gene symbols using biomaRt package (v2.44.4) with Ensembl as the host. We used our 2 DEG lists (c.4596_4597insG vs. WT; c.4582-2delAG vs. WT) and investigated overlapping genes (up/down/both) separately. Genes with adjusted p values <0.05 and absolute log_2_FC > 0 were called DEGs.

### SynGo annotation

For analysis of DEGs associated with synaptic function, GO analysis was performed using the SynGO [55] website: https://www.syngoportal.org/.

### WGCNA

Gene counts were normalized using ComBat seq ^33^ to remove the batch effect. The network was constructed using the WGCNA package (v1.70.3) ^34^ through a signed network. A soft-threshold power of 18 was used to achieve approximate scale-free topology (R^2^>0.8). Module GO term enrichment was performed using cluster Profiler (v4.0.5)^35^ with default parameters. The module network is visualized using Cytoscape (v 3.10.2)^36^.

### eSNP-Karyotyping analysis

#### RNA-Seq data processing

RNA-Seq data were initially processed using the nf-core/rnaseq pipeline (version 3.12.0) executed via Nextflow (version 23.04.4)^37^. In detail, we merged the raw FastQ files using the *cat* (version 8.30) command to consolidate data from technical replicates. Next, we auto inferred the strandedness of the libraries through subsampling and pseudo-alignment using *fq* (version 0.9.1 (2022-02-22) and *Salmon* (version 1.10.1)^38^. Then, we conducted quality control checks with FastQC (version 0.11.9), and trimmed adapters and low-quality sequences using Trim Galore! (version 0.6.7).

After we obtained the trimmed reads, we combined trimmed paired-end reads and aligned them to the human reference genome (GRCh38) using the *Bowtie2* aligner (version 2.4.1), resulting in a set of Sequence Alignment/Map (SAM) format files [Bowtie2]. We then converted the SAM file to binary alignment and map (BAM) format for downstream processing using *Samtools* View (version 1.3.1) with the “-bS” flag enabled^39^.

#### Cell line karyotyping

First, we called variants from pre-processed BAM files against the reference genome using the GPU-accelerated *haplotypecaller* available with the *NVIDIA Parabricks toolkit* (version 4.3.1, https://docs.nvidia.com/clara/parabricks/4.3.1/).

We then assessed the potential presence of chromosomal multiplications using *eSNP-karyotyping* method^40^, analyzing the binary alignment map (BAM) and VCF files in conjunction with dbSNP (version 155)^41^. Following previously published methods, we compared the allelic ratio in sliding windows of 151 single-nucleotide polymorphisms (SNPs) with a minimal allele frequency above 0.2 and were covered by more than 20 reads with the rest of the genome. We identified significant multiplications as those with a false discovery rate below 0.05.

### Quantification and statistical analysis

All data are presented as the mean ± standard error of the mean (SEM), unless otherwise noted. All experiments were repeated on at least two-three biological replicates for each isogenic pair, with each replicate consisting of an independent infection/differentiation. Statistical significance ((*p < 0.05, **p < 0.01; ***p < 0.001) was evaluated with the Kolmogorov–Smirnov-test (for cumulative probability plots), two-way ANOVA, and Student’s t-test (for bar-graph).

## Results

### Generation of a *SETD1A^+/-^* LoF iPSC lines mimicking patient with *SETD1A* haploinsufficiency (c.4582-2delAG and c.4596_4597insG (p. Leu1533fs))

To study patient-specific mutations in an iPSC-derived human neural model, we generated five pairs of isogenic *SETD1A^+/-^*LoF mutation iPSC lines using the CRISPR-Cas9 gene editing technology (**Fig 1A**). First, we generated a patient-specific knock-in mutation c.4582-2delAG by targeting the Intron 15-Exon 16 boundary of *SETD1A* in two healthy control iPSC lines (CD07 and C11C). Meanwhile, we also created an exon 16 frameshift (fs) mutation by generating a 1-base pair (bp)-insertion mutation c.4596_4597insG (p. Leu1533fs) in three healthy control iPSC lines (CD07, CD15, CD19) to mimic *SETD1A^+/-^* LoF caused by c.4582-2delAG. The mutated DNA sequences were confirmed by Sanger sequencing (**Fig 1A**), and no off-targets were found for the top 5 predicted loci (**Table 1**). The c.4582-2delAG in *SETD1A* disrupted the splicing adaptor sequences and retained the intron 15, which led to a short fragment of *de novo* protein sequence after exon 15 and a premature stop codon in exon 16 (**Fig 1A&B**). The c.4596_4597insG in exon 16 led to a premature stop codon and thus, protein truncation. Both mutations led to premature stop codons and are predicted to disrupt the SET domain function, the key to the histone methyltransferase function. These gene targeting strategies did not yield iPSC lines carrying homozygous *SETD1A* mutations, which is consistent with the literature that shows *SETD1A* homozygous knockouts in germline cells as lethal ^42^. As a result, five pairs of isogenic heterozygous *SETD1A* LoF mutation iPSCs were generated to model humans with heterozygous *SETD1A* LoF alleles. The pluripotency of all generated iPSC lines was confirmed through immunostaining with pluripotency markers OCT4, SOX2, and Tra1-60 (**Fig S1A**).

**Fig 1.**
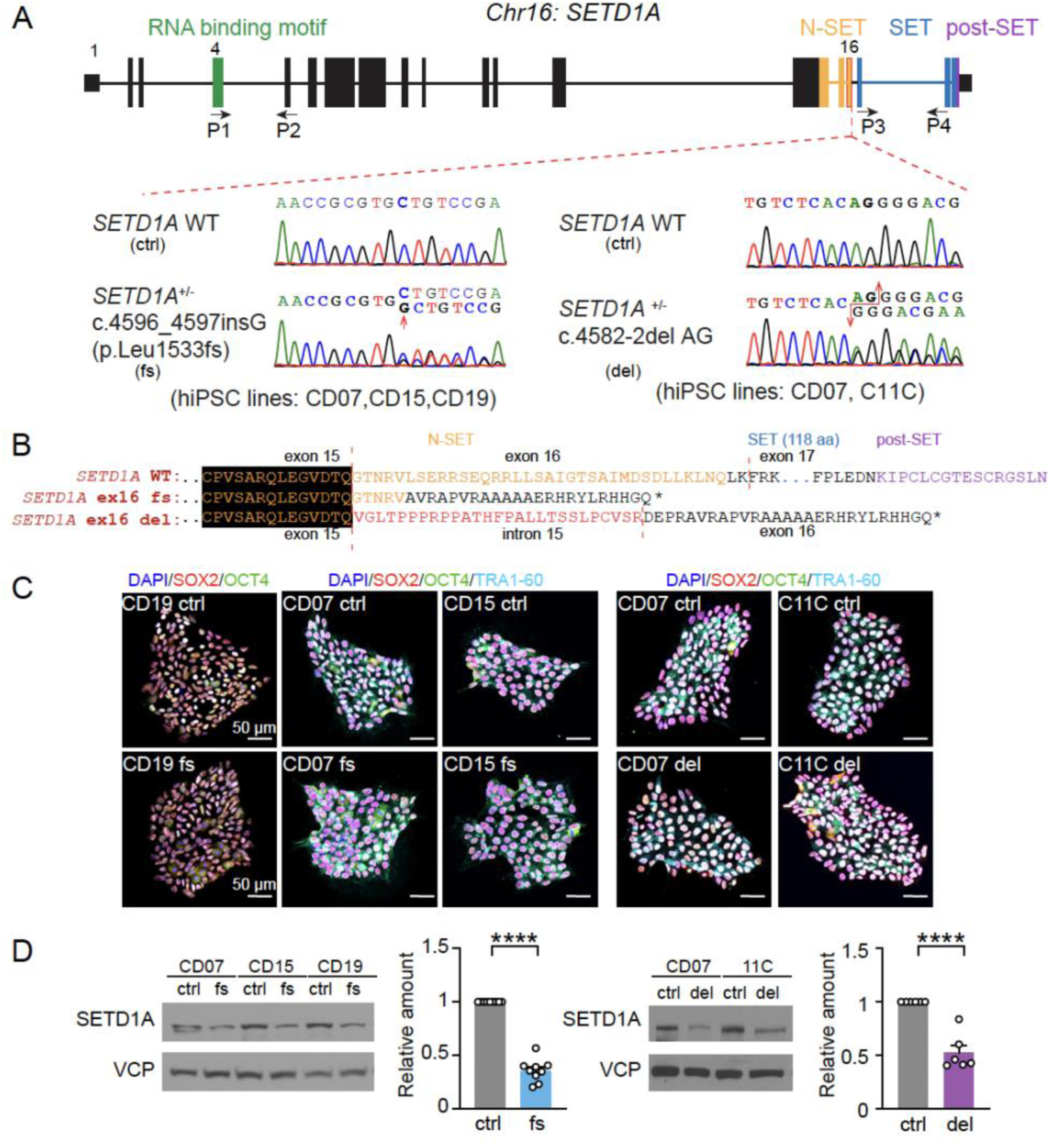
Generation and validation of iPSCs carrying *SETD1A*^+/-^ *LoF* mutations. (**A**) Diagram showing the genomic structure of *SETD1A* and the two sets of mutations. Two sets of *SETD1A*^+/-^ LoF mutations (c.4596_4597insG (p.Leu1533fs) and c.4582-2del AG) in four different genetic backgrounds (CD07, CD15, CD19, C11C) were generated. *Low panels:* Sanger sequencing confirms the mutations. (**B**) The protein sequence of SETD1A and predicted protein sequence of *SETD1A*^+/-^ LoF mutations. (**C**) Representative images of iPSCs with pluripotency markers SOX2, OCT4 and TRA1-60 and nucleus marker DAPI. Scale bars: 50μm. (**D**) Western blot of SETD1A protein normalized to VCP in iPSCs. N=9 for c.4596_4597insG mutation (3 independent batches of 3 isogenic pairs) and N= 4 for c.4582-2del AG mutation (2 independent batches of 2 isogenic pairs). Statistical significance was assessed by unpaired t-test. Data are shown as means ± SEM. *P < 0.05, **P < 0.01, ***P < 0.001 and **** P < 0.0001.

We next examined the expression of SETD1A protein to confirm the LoF premature stop codon-mutations in the *SETD1A* gene using Western blotting. Consistent with the reported data from the isolated patients’ lymphoblastoid cell lines carrying the same mutation (c.4582-2delAG) ^20^, we observed a significant decrease of full-length SETD1A protein levels in iPSCs carrying c.4582-2delAG and c.4596_4597insG mutations through two sets of SETD1A antibodies that target either exon 2 or exon 14 of *SETD1A*, respectively (**Fig 1D, Fig S2A**).

### Mutant *SETD1A* mRNAs are degraded through nonsense-mediated decay (NMD)

Having demonstrated that iPSCs carrying c.4582-2delAG and c.4596_4597insG mutations cause a reduction in SETD1A protein expression (**Figs 1D, S2A**), we hypothesize that mutant *SETD1A* mRNAs are degraded. Thus, we quantified the *SETD1A* mRNA level and tested the mechanisms that regulate *SETD1A* mutant mRNA degradation. It is known that NMD is a translation-dependent mRNA surveillance pathway that reduces errors in gene expression by eliminating mRNA transcripts that contain premature stop codons ^43^. Although the SCZ patient-specific mutations c.4582-2delAG and c.4596_4597insG are both predicted to generate premature-stop codon in exon 16, it is unknown whether these mutations indeed cause NMD of *SETD1A* mRNA. Towards this end, we first tested whether the mRNA was degraded by using two different sets of primers that target different transcript regions before or after the CRISPR-introduced mutations (P1+P2 across exons 4-5 and P3+P4 for exons 17-18). Consistent with the ∼50% reduction of full-length SETD1A protein level in iPSCs (**Fig 1D**), we found ∼50% reduction of *SETD1A* mRNA level in iPSCs when interrogating the two different transcript regions (**Fig 2A**), suggesting NMD may occur for the mRNAs produced by the LoF alleles (c.4582-2delAG and c.4596_4597insG).

**Fig. 2.**
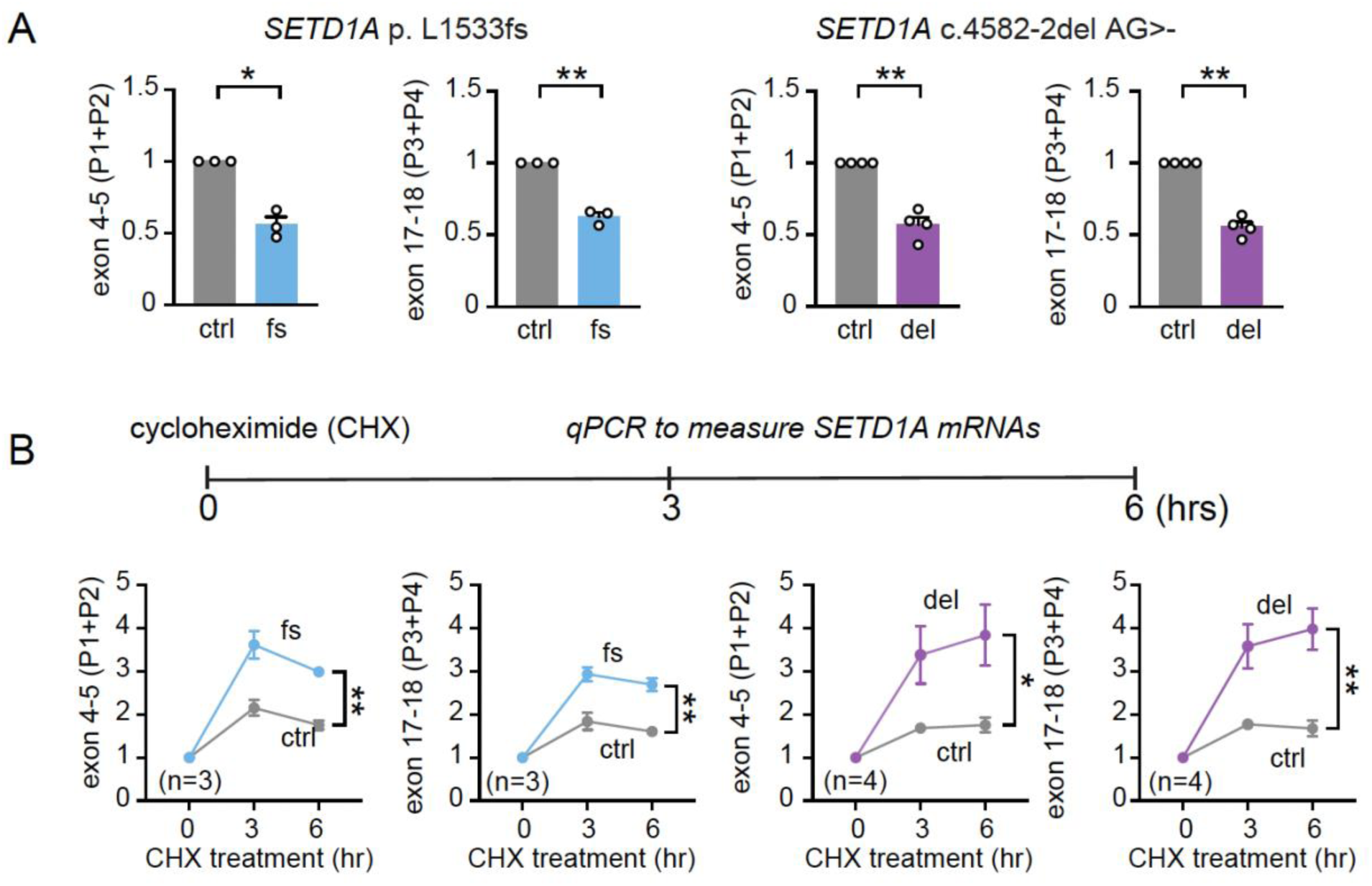
Down-regulation of mRNAs carrying *SETD1A*^+/-^ LoF is mediated by NMD. (**A**) Normalized *SETD1A* mRNA expression in *SETD1A^+/-^* LoF iPSCs using two sets of primer as depicted in Figure 1A: P1+P2 for exon 4-5 and P3+P4 for exon 17-18. (**B**) Impact of cycloheximide (CHX) exposure (0, 3, 6 hours) on *SETD1A* mRNA expression levels in iPSCs carrying *SETD1A^+/-^* LoF. N= 3 for c.4596_4597insG mutation (1 batch of 3 isogenic pairs) and N= 4 for c.4582-2del AG mutation (2 batches of 2 isogenic pairs). Statistical significance was assessed by t-test for qPCR in **A** and two-way ANOVA for qPCR after CHX treatment in **B**. Data are shown as means ± SEM. *P < 0.05, **P < 0.01, ***P < 0.001 and **** P < 0.0001.

To further corroborate whether these *SETD1A* mutations indeed lead to NMD of the mRNA, we treated iPSCs with a translation inhibitor cycloheximide (CHX,100ug/ml) for 3 and 6 hours and then examined the expression level of *SETD1A* mRNA. CHX inhibits protein translation and subsequently interferes with the NMD machinery’s recognition of a premature stop codon. We observed that *SETD1A* transcripts increased dramatically after CHX treatment in iPSC lines carrying c.4596_4597insG or c.4582-2delAG mutation when compared with control wild-type (WT) iPSCs (**Fig 2B**). These results indicate that both patient-specific mutations c.4582-2delAG and c.4596_4597insG in exon 16 of *SETD1A* cause protein LoF through NMD of *SETD1A* mRNAs.

### *SETD1A*^+/-^ LoF excitatory Ngn2-iNs showed decreased dendrite complexity

To investigate the impact of *SETD1A*^+/-^ LoF on the human neural model, we generated excitatory Ngn2-iNs (**Fig 3A**) using both control and mutant iPSCs. These iNs were replated on monolayer mouse astrocytes to facilitate their maturation and synapse formation ^44^. Mutant iPSCs carrying *SETD1A*^+/-^ LoF mutations, i.e., c.4582-2delAG or c.4596_4597insG, could produce mature Ngn2-iN cells similar to control iPSCs. We next examined the expression of SETD1A proteins by Western blotting. Similar to that found in iPSC lines, we observed ∼50% reduction of full-length SETD1A proteins when compared with control neurons (**Fig 3B&C**).

**Fig. 3.**
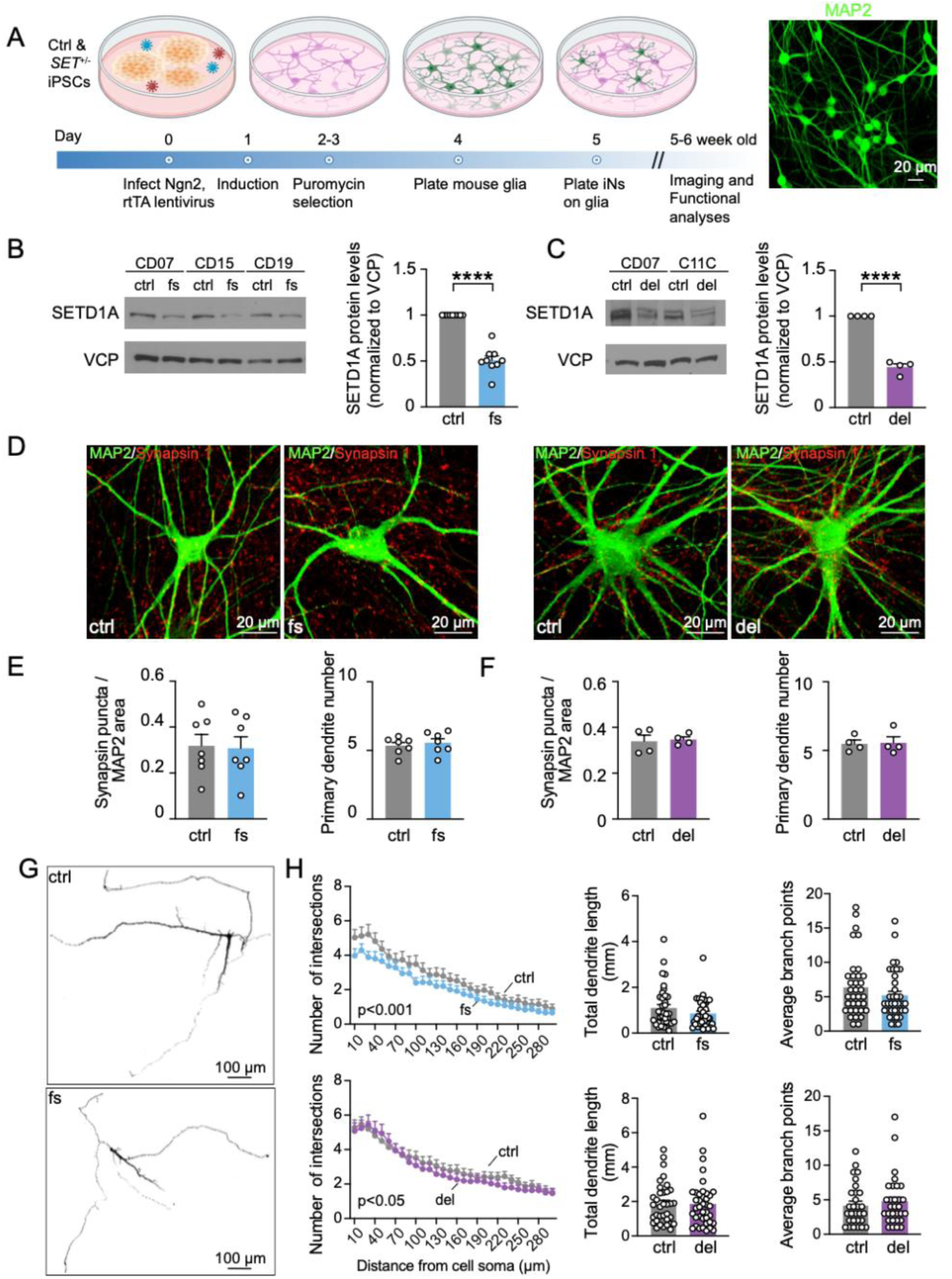
Morphological characterization of Ngn2-iNs carrying *SETD1A*^+/-^ LoF mutations. (**A**) Diagram showing the experimental design and the timeline of Ngn2-iNs culture. (**B&C**) Western blot of normalized SETD1A protein expression in Ngn2-iNs for c.4596_4597insG and c.4582-2delAG mutations, respectively. N=9 for c.4596_4597insG mutation (3 independent batches of 3 isogenic pairs) and N= 4 for c.4582-2del AG mutation (2 independent batches of 2 isogenic pairs). (**D**) Representative images of Ngn2-iNs with IHCs of neuronal marker MAP2 and pre-synaptic marker synapsin-1 in control and *SETD1A^+/-^* mutant neurons. (**E&F**) Quantification of synaptic puncta number normalized to MAP2-labelled dendrites and the number of primary dendrites for c.4596_4597insG and c.4582-2del AG mutations, respectively. Each dot represents the average of one batch of one isogenic pair. N= 7 for c.4596_4597insG mutation (3 batches of 1 isogenic pair and 2 batches of 2 isogenic pairs). N= 4 for c.4582-2delAG mutations (2 batches of 2 isogenic pairs). (**G**) Representative images of neurons sparsely labeled by GFP for dendrite morphology analysis. (**H**) Quantification of neuronal dendrite morphology using *Sholl* analysis. Total dendrite length and average branch points were quantified. Each dot represents one neuron in bar graph. N= 34 for control and N= 36 for c.4596_4597insG from 3 independent batches. Statistical significance was assessed by unpaired t-test (bar graphs) and two-way ANOVA (*Sholl* analysis). Data are shown as means ± SEM. *P < 0.05, **P < 0.01, ***P < 0.001 and **** P < 0.0001.

We next examined the impact of *SETD1A*^+/-^ LoF mutations on synaptogenesis. We conducted immunostaining with dendritic marker MAP2 and presynaptic marker synapsin-1 (**Fig 3D**) for 5 to 6-week-old excitatory human iNs. We used a custom-made algorithm, *Intellicount* ^30^, to unbiasedly quantify synapsin-1 labeled synaptic puncta colocalized with MAP2-labelled dendrites and found no significant differences between control and mutant neurons (**Fig 3E&F**).

To further characterize the impact of *SETD1A*^+/-^ mutations on the morphogenesis of human neurons, particularly the neuronal dendrite development, we sparsely labeled neurons with pMax-GFP. We reconstructed neuronal 3D structure to analyze the dendrite complexity and length (**Fig 3G**). We found that mutant neurons carrying the c.4582-2delAG and c.4596_4597insG mutations displayed reduced dendrite complexity through *Sholl* analysis without affecting the dendrite length and number of branch points (**Fig 3H**).

### *SETD1A*^+/-^ LoF alters synaptic transmission and short-term synaptic plasticity in Ngn2-iN cells

To investigate the impact of *SETD1A*^+/-^ LoF on neuronal functions, including intrinsic electrophysiological properties and synaptic transmission, we performed whole-cell patch clamp recordings from 5 to 6-week-old excitatory human iNs cultured on monolayer astrocytes. We first measured intrinsic electrophysiological parameters, such as cell membrane capacitance and resistance (**Fig S3A-D**). We observed increased membrane capacitance and decreased membrane resistance for c.4596_4597insG but no significant differences for c.4582-2delAG. Next, we characterized the neuronal excitability and analyzed the action potential (AP) kinetics (**Fig S4**). We found that neurons carrying c.4582-2delAG mutation showed increased neuronal excitability without affecting the kinetics of APs, including AP threshold, amplitude, afterhyperpolarization potential (AHP), and half-width of APs.

Synaptic transmission is vital for neuronal communication and signal transmission. Next, we asked whether *SETD1A*^+/-^ LoF mutations alter synaptic function in human iNs. We first analyzed miniature excitatory postsynaptic currents (mEPSCs) (**Fig 4**). We found that mutant iN cells showed higher mEPSC frequency for both c.4596_4597insG and c.4582-2delAG mutations (**Fig 4B&E**). However, we did not observe significant differences for mEPSC amplitude except for altered cumulative probability distribution of amplitudes in neurons carrying c.4582-2delAG mutation (**Fig 4C&F**). These data suggest that the overall spontaneous miniature synaptic transmission is enhanced in human neuronal cells carrying *SETD1A^+/-^* LoF mutations.

**Fig. 4.**
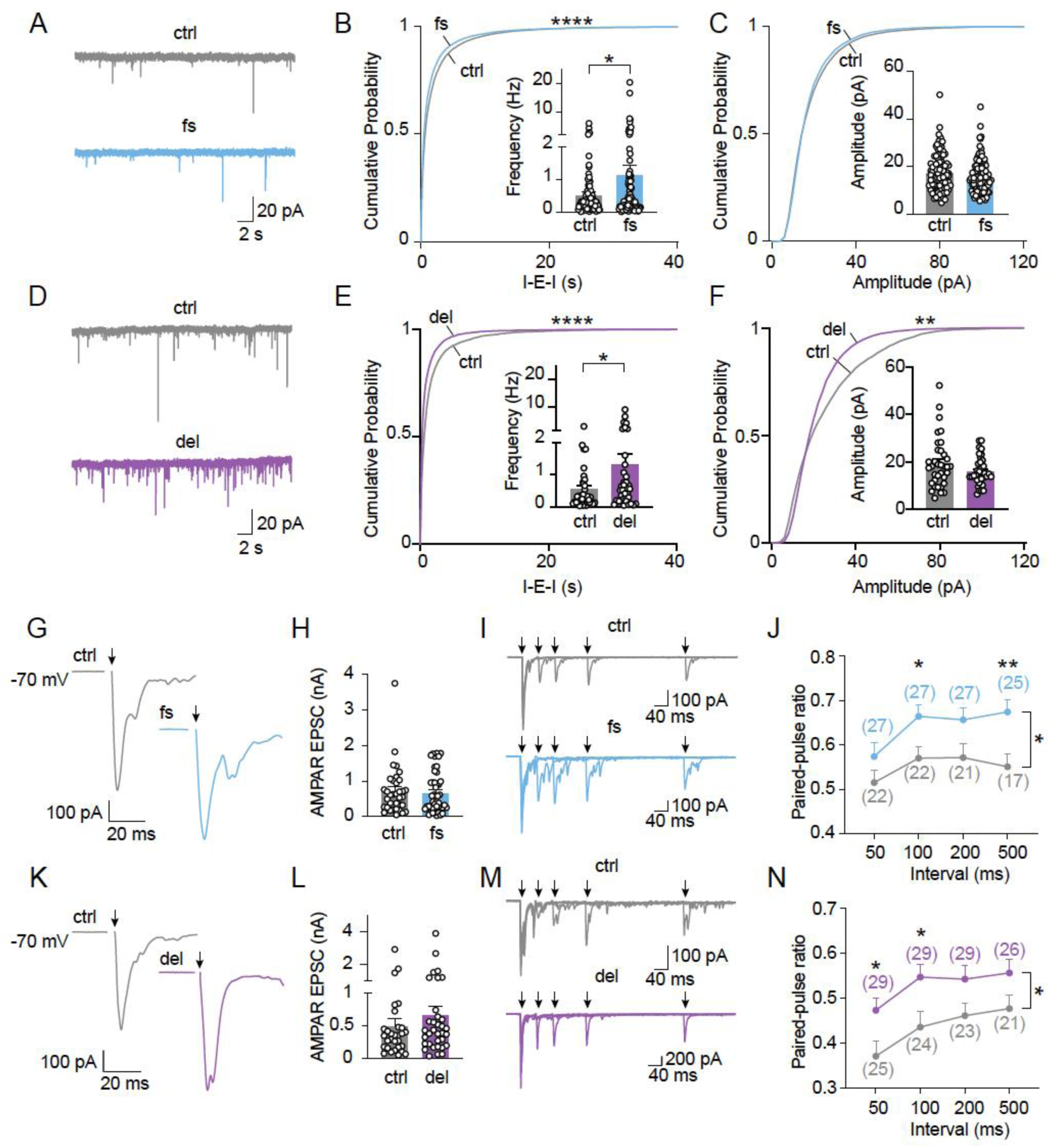
Functional characterization of Ngn2-iNs carrying *SETD1A*^+/-^ LoF mutations. (**A**) Representative traces of mEPSC recording for control and c.4596_4597insG mutant Ngn2-iNs. (**B&C**) Cumulative distribution and quantification of mEPSC frequency (inter-event-interval, I-E-I) in (**B**) and mEPSC amplitude in (**C**). (**D**) Representative traces of mEPSC recording for control and c.4582-2delAG mutant Ngn2-iNs. (**E&F**) Cumulative distribution and quantification of mEPSC frequency (inter-event-interval, I-E-I) in (**E**) and mEPSC amplitude in (**F**). (**G**) Representative traces of evoked AMPAR EPSC for control and c.4596_4597insG mutant Ngn2-iNs. (**H**) Quantification of evoked AMPAR EPSC amplitude. (**I**) Representative traces of paired-pulse recording traces for control and c.4596_4597insG mutant Ngn2-iNs. (**J**) Quantification of paired-pulse ratio. (**K**) Representative traces of evoked AMPAR EPSC for control and c.4582-2delAG mutant Ngn2-iNs. (**L**) Quantification of evoked AMPAR EPSC amplitude. (**M**) Representative traces of paired-pulse recording traces for control and c.4582-2delAG mutant Ngn2-iNs. (**N**) Quantification of paired-pulse ratio (PPR). Statistical significance was evaluated with the Kolmogorov–Smirnov test (cumulative probability plots), unpaired t-test (frequency and amplitude) and two-way ANOVA (paired-pulse ratio plots). *P < 0.05, **P < 0.01, ***P < 0.001 and **** P < 0.0001.

To further investigate the synaptic alterations by *SETD1A^+/-^*LoF mutations, we recorded evoked EPSCs (eEPSCs) by an extracellular stimulation electrode ^45^ in mutant *SETD1A^+/-^* LoF iNs. We did not observe changes in evoked EPSCs amplitude in either c.4596_4597insG or c.4582-2delAG mutant *SETD1A*^+/-^ LoF iNs (**Fig 4G&K**). We then assayed the short-term synaptic plasticity using a paired-pulse protocol with intervals between 50 to 500 ms. Because of no apparent differences in the sizes of eEPSCs between the control and the mutant iNs, we were expecting no change in the paired-pulse ratio (eEPSC2/eEPSC1). To our surprise, we observed a paradoxical, but consistent, increase in the PPRs, suggesting an alteration in short-term synaptic plasticity (recovering or replenishment of readily releasable pool of synaptic vesicles) in neurons carrying *SETD1A*^+/-^ LoF mutations (**Fig 4I-N)**.

Collectively, our electrophysiological data suggest that neurons with patient-specific *SETD1A*^+/-^ LoF mutations c.4582-2delAG and c.4596_4597insG exhibited altered neuronal excitability, dysregulated synaptic transmission, and short-term synaptic plasticity.

### Transcriptional profiling characterization of *SETD1A^+/-^* LoF Ngn2 iNs

To understand how *SETD1A*^+/-^ LoF mutations (c.4596_4597insG and c.4582-2delAG) altered synaptic function in human iN cells, we conducted a transcriptomic analysis using bulk RNA sequencing (RNA seq) in 6-week-old iNs of both control and mutant neurons. Our RNA seq results confirmed the absence of chromosomal abnormality in all 5 isogenic pairs (**Fig S5**) by eSNP-Karyotyping. We then examined the expression level of *SETD1A*. Consistent with our qPCR data (**Fig 2A**), we found that *SETD1A* is significantly downregulated in *SETD1A*^+/-^ LoF mutant human iN cells (**Figs 5A&B, S7A**). Along with the changes of *SETD1A*, we identified a substantial number of differentially expressed genes (DEGs) (493 DEGs for c.4596_4597insG and 1705 DEGs for c.4582-2delAG) in *SETD1A* mutant neurons when compared with control human iN cells (**Fig 5A&B**). The expression levels of some presynaptic genes, such as *DOC2B* (**Fig S7B**), which is involved in regulating spontaneous synaptic releases ^46^, are significantly increased in iNs carrying both c.4596_4597insG and c.4582-2delAG mutations.

**Fig. 5.**
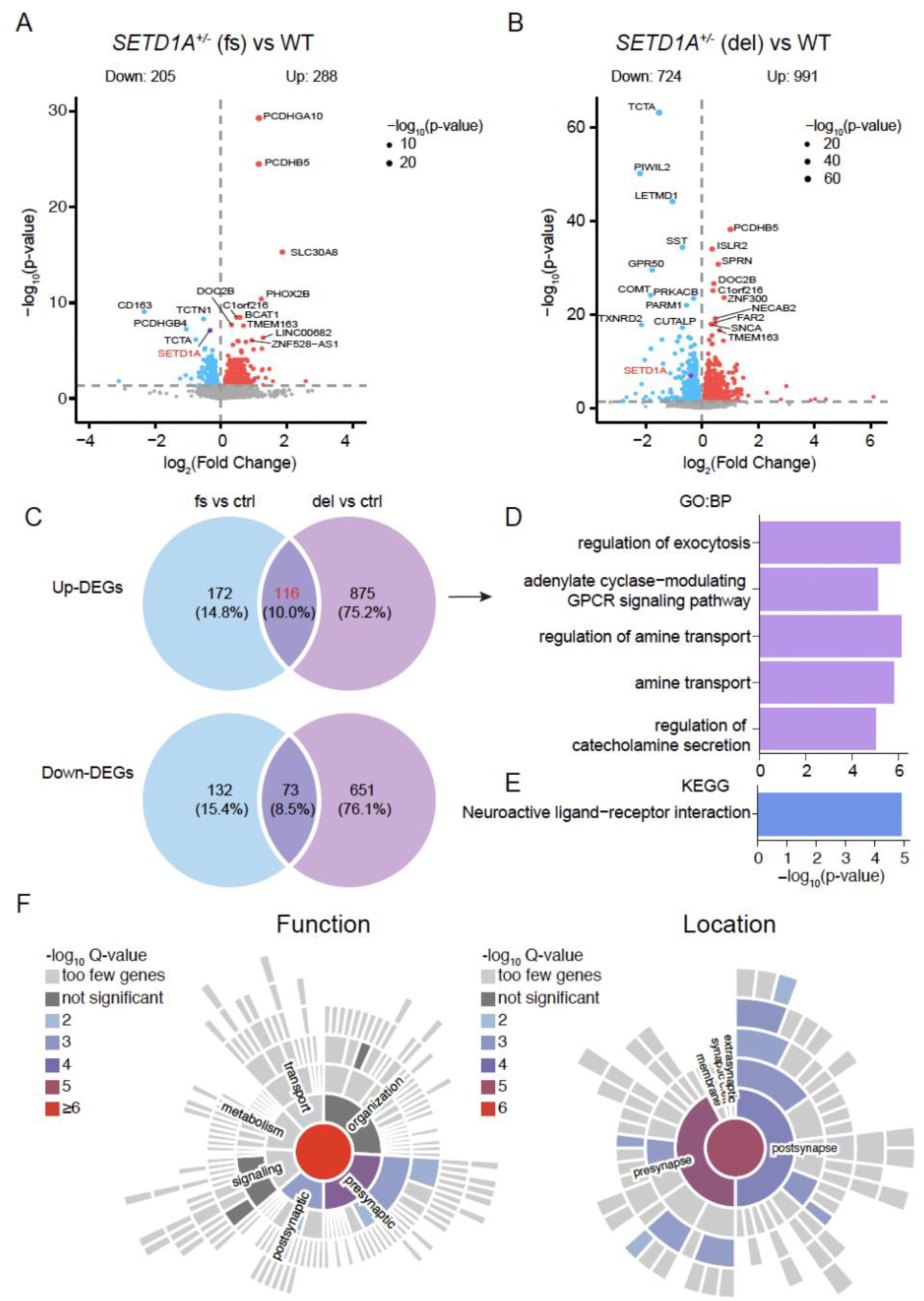
Transcriptomic analysis of Ngn2-iNs carrying *SETD1A*^+/-^ LoF mutations. (**A&B**) Volcano plot of Ngn2-iNs carrying c.4596_4597insG and c.4582-2delAG mutations compared to control. Differential expression is defined by adjusted p-value < 0.05 and log_2_FC > 0. (**C**) Venn diagram shows the overlapped DEGs between c.4582-2delAG and c.4582-2delAG mutations. (**D-F**) The enriched GO biological pathways(**D**), enriched KEGG terms (**E**) and Syn-go function and location enrichment analysis (**F**) of overlapped 115 up-regulated DEGs.

To understand the biological pathways and functions of the dysregulated DEGs, we conducted Gene Ontology (GO) enrichment analyses. For *SETD1A* c.4596_4597insG mutation, the up-regulated genes (n=288) showed significant enrichment for synaptic function and amine transport-associated pathways; however, the down-regulated genes (n=205) did not show any significantly enriched pathways (**Fig S6A**). For *SETD1A* c.4582-2delAG mutation, the up-regulated genes (n=991) are also enriched in synaptic function and amine transport-associated biological pathways (**Fig S6B**). In addition, the down-regulated genes (n=724) are also enriched for axon development and axon guidance-related pathways (**Fig S6C**). Interestingly, we also found that several metabolic pathways, such as glycolysis and Adenosine Diphosphate (ADP) metabolic process, are enriched in the up-regulated DEGs in c.4582-2delAG dataset, suggesting a wide-spread effect of *SETD1A^+/-^* LoF mutation that may be beyond direct neuron-related function.

Given the similar effects of the patient-specific mutation c.4582-2delAG and the other exon-16 mutation c.4596_4597insG that mimics c.4582-2delAG, we further narrowed down the DEGs by examining the shared DEGs in these two datasets (**Fig 5C**). We identified 116 shared up-regulated DEGs and 73 shared down-regulated DEGs (**Fig 5C**). Consistent with previous GO analysis results, GO enrichment analysis revealed that the overlapped up-regulated genes are enriched for biological pathways related to regulation of exocytosis and amine transport (**Fig 5D&E**), which highlights the impact of *SETD1A*^+/-^ LoF on synaptic function. To further understand the function of these overlapped 116 up-regulated DEGs, we did Synaptic Gene Ontologies (Syn-GO) analysis ^47^ and found genes associated with synaptic function significantly are enriched in the presynaptic terminal (**Fig 5F**). To further gain a gene network view of these DEGs, we conducted protein-protein-interaction (PPI) analysis using Search Tool for the Retrieval of Interacting Genes (STRING) ^48^. The PPI network of 116 overlapped up-regulated DEGs was constructed with a medium confidence score (0.4) based on both functional and physical protein associations. We found significant enrichments of several biological processes, including regulation of amine transport, postsynaptic membrane potential, chemical transmission, and exocytosis (**Fig S7C**). The PPI analysis of cellular component showed proteins are enriched mainly in the axon terminus, pre-, and post-synaptic membrane (**Fig S7D**). These results largely support that dendritic arborization is altered in *SETD1A^+/-^* LoF human iN cells, and synaptic transmission is enhanced (mEPSCs frequency and recovery of the readily releasable pool of the synaptic vesicles) (**Fig 4B&D**).

To further understand the regulated transcriptional network, we constructed gene networks to identify modules of highly co-expressed genes, a powerful method to identify biological processes in an unsupervised manner ^49,50^. We conducted weighted gene co-expression network analysis (WGCNA) analysis ^34^ by combining the c.4596_4597insG and c.4582-2delAG mutation datasets together. Here, we identified 20 modules in total with seven of them showing a significant correlation with *SETD1A*^+/-^ LoF mutation (**Fig 6A**). To understand the biological pathways enriched in these seven modules, we conducted GO analysis for genes in each module (**Fig 6B**). We found that six out of the seven *SETD1A*^+/-^ LoF mutation-associated modules showed significant enrichment for biological pathways, such as metabolic pathways (M19 and M12), development of nervous system (M4), synaptic function and membrane potential regulation (M10), extracellular matrix organization (M20), and skeletal system morphogenesis (M15) (**Fig 6B**). In addition, gene co-expression networks of 6 *SETD1A*^+/-^ LoF mutation-associated modules are visualized using Cytoscape ^36^ (**Fig 6C**). Interestingly, *SETD1A* was included in M15, a module showing significant association with *SETD1A*^+/-^ LoF mutation, indicating *SETD1A* as a critical regulator of gene co-expressions in M15. These results suggest that human iNs carrying the *SETD1A*^+/-^ LoF mutations have altered gene co-expression networks that are mainly enriched in cell metabolism, synaptic function, and development of nervous system, underscoring the specific impact of *SETD1A*^+/-^ LoF on human neuronal function, metabolism, and development.

**Fig. 6.**
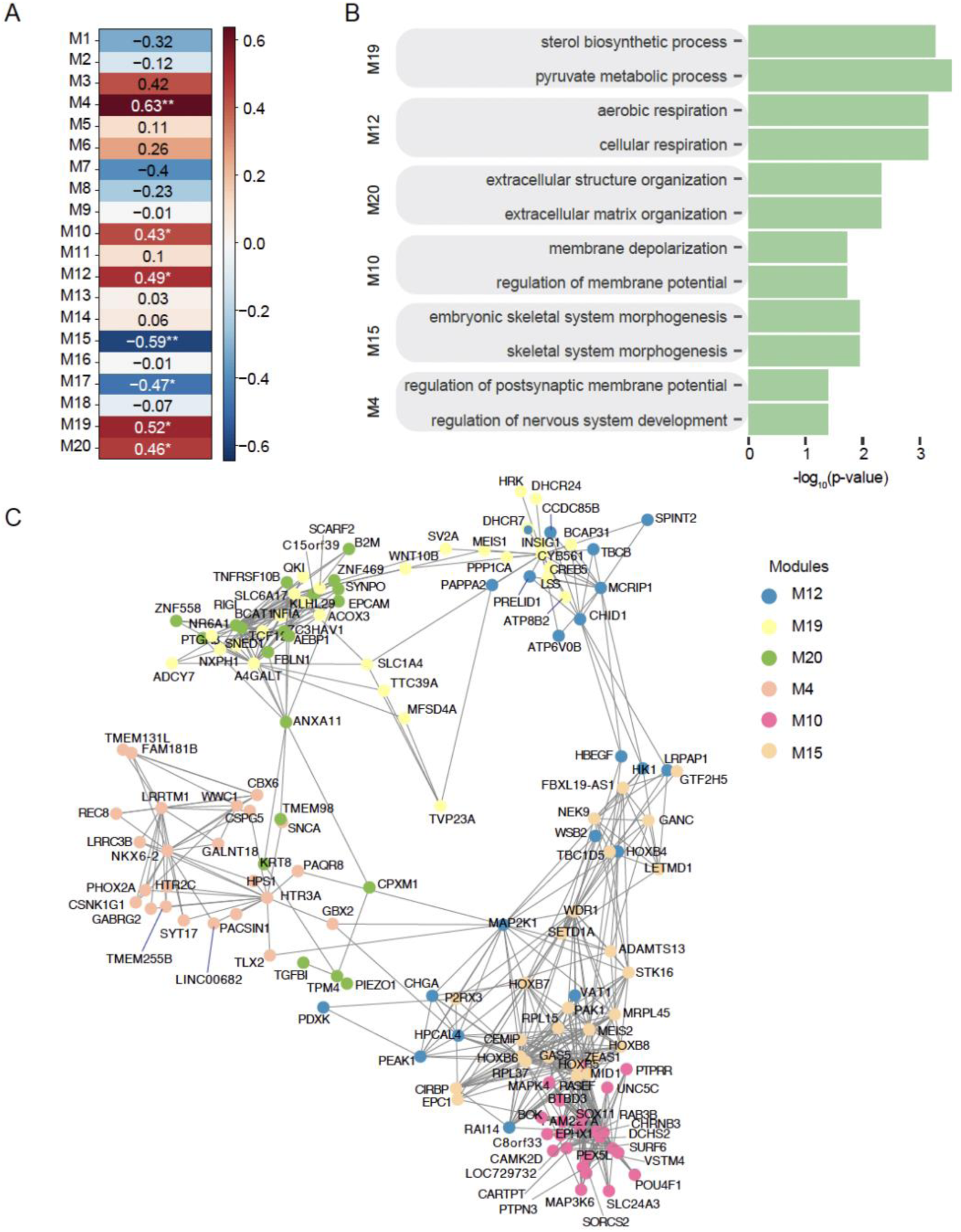
WGCNA analysis of Ngn2-iNs carrying *SETD1A*^+/-^ LoF mutations. (**A**) Heatmap shows the correlation between the identified 20 modules and phenotype (*SETD1A^+/-^* LoF mutations). (**B**) The enriched GO biological pathways for modules are significantly associated with *SETD1A^+/-^* LoF mutations. (**C**) The gene co-expression networks in 6 modules are visualized by Cytoscape.

## Discussion

Using human neuronal cells derived from five isogenic pairs of iPSCs carrying the patient-specific c.4582-2delAG and c.4596_4597insG (p.Leu1533fs) mutations in *SETD1A* gene, we investigated the impact of *SETD1A*^+/-^ LoF on neuronal transcriptional profiling, morphology, and functions. We found that the patient mutation c.4582-2delAG in *SETD1A* causes *SETD1A* haploinsufficiency and dysregulates synapse functionality. We found that both c.4582-2delAG and c.4596_4597insG cause a reduction of SETD1A full-length protein expression, which is accompanied by a reduction in *SETD1A* mRNA expression levels. Morphological analysis revealed that *SETD1A*^+/-^ LoF affects dendritic development; electrophysiological analysis revealed an altered synaptic transmission in human iNs carrying *SETD1A*^+/-^ LoF mutations; transcriptional analysis identified DEGs that are associated with dendritic morphogenesis and synaptic transmission. Our findings indicate that patient-specific *SETD1A* mutation (i.e., c.4582-2delAG) dysregulates synaptic function.

Large-scale human genetics studies in SCZ have identified that SCZ cases carry a significantly higher burden of rare PTVs among evolutionarily constrained genes across different ancestries ^2,51^. The interpretation of the effects of the rare PTVs is important for the accurate determination of variant impacts. It has been shown that NMD effects are highly concordant across tissues and individuals, which may at least partially address the impact of PTVs in humans ^52^.

Interindividual differences of RNA stability may account for the heterogenous phenotypical expression of the disease phenotype in humans ^53^. Based on previous findings, however, SCZ-associated *OTUD7A* LoF causes cellular phenotype without NMD in human neurons ^54^, suggesting that NMD is not the only hypothesis related to SCZ PTVs. Our analyses indicated that both c.4582-2delAG and c.4596_4597insG mutations lead to the reduction of *SETD1A* mRNA and protein levels. The patient-specific c.4582-2delAG mutation is associated not only with SCZ but also NDDs ^7^. Our results indicate that *SETD1A* mRNA reduction can be restored directly through inhibiting NMD by CHX in human iPSCs. It is highly suggested that NMD plays a role in regulating *SETD1A* mRNAs and protein expressions in humans. Since there are many different types of mutations identified in *SETD1A* ^7^, further studies are required to precisely interpret different mutations accounting for the heterogeneous genetic causes.

Our morphological results demonstrate that *SETD1A*^+/-^ LoF mutations decrease dendrite complexity without affecting synapse formation and dendrite length. This is largely consistent with previous studies using both mouse and human neuron culture models for *SETD1A*^+/-^ LoF ^17–19,21,22^. Three studies of *Setd1a* haploinsufficiency in mouse models revealed deficits in dendrite development and decreased dendritic spine density ^17–19^. However, human neuronal studies about excitatory and inhibitory neurons in vitro showed increased dendrite length at a later time point without synaptic puncta formation changes ^21^. Another cortical spheroid model showed decreased neurite outgrowth in cortical neurons ^22^. Overall, both in vivo and in vitro studies converge in altered dendrite complexity as a common morphological phenotype for *SETD1A*^+/-^ LoF. Although the detailed mechanism remains unclear, our transcriptomic studies revealed that the expression of genes associated with axon development is downregulated by *SETD1A*^+/-^ LoF.

The synaptic hypothesis of SCZ has been proposed to be the underlying convergent biological pathway that contributes to the pathophysiology of SCZ ^55^. Recent GWAS studies also support synaptic dysregulation as a major biological process associated with SCZ and other NDDs ^3^. Using human neurons as a model system, our study demonstrates that *SETD1A*^+/-^ LoF mutations lead to synaptic dysregulation, including enhanced synaptic transmission, enhanced neuronal excitability, and altered synaptic plasticity. The increased miniature excitatory synaptic transmission and enhanced neuronal excitability suggest hyperactivity in excitatory neurons and possible increased release probability in the presynaptic compartments resulting from *SETD1A*^+/-^ LoF. However, our evoked recording results indicate that there is no difference in AMPAR-mediated EPSC and significantly increased PPR in neurons carrying *SETD1A*^+/-^ LoF, suggesting potential decreased releasing probability in presynaptic terminals. This apparent paradoxical phenotype could be explained by an increased recovery of readily releasable pools after each release in the *SETD1A* mutant human neurons. Interestingly, a presynaptic protein DOC2B showed a consistent increase in mRNA expression levels from the RNA seq data of the *SETD1A* mutant human neurons (**Fig S7B**). It has been shown that DOC2B is an important modulator for spontaneous synaptic releases and asynchronous synaptic release ^46,56–58^. The knockdown of DOC2 families leads to decreased frequency of spontaneous release without affecting evoked release, and it appears that DOC2 protein in the presynaptic compartment regulates spontaneous release in a Ca^2+^-independent manner ^46^. Although the detailed mechanisms remain to be elucidated, the observed altered synaptic transmission and synaptic plasticity are also supported by mouse prefrontal cortex slice recording ^17,18^, indicating synaptic dysregulation as a common phenotype caused by *SETD1A*^+/-^ LoF.

Transcriptomic analysis reveals up-regulation of synaptic transmission, ion transport, and cell metabolism-related biological pathways and down-regulation of axon development and guidance. These data support an enhanced synaptic function and defective dendritic development in human neurons that we found in this study. This is also consistent with the previously reported gene expression data in both mouse and human neuron studies ^17,21^. When the DEGs from the *SETD1A*^+/-^ LoF neurons were compared with DEGs with the chromatin immunoprecipitation sequencing (ChIP-seq) database of SETD1A from mouse prefrontal cortex ^17^, we found 58 overlapping DEGs and 18 overlapping DEGs for c.4582-2delAG and c.4596_4597insG (p.Leu1533fs), respectively.

This study on *SETD1A*^+/-^ LoF in the human neuronal system also has limitations. It mainly focuses on studying the impact of *SETD1A*^+/-^ LoF in induced human excitatory neurons, which transdifferentiates iPSCs to induced excitatory neurons by ectopic expression of transcription factor Ngn2 ^23^. This model may not fully capture the differentiation progress that occurs during the neurodevelopmental stage, as our study solely focuses on excitatory neurons. It’s known that SETD1A is broadly expressed in all brain cell types, including other neuronal subtypes such as inhibitory neurons, dopaminergic neurons, and glial cells. Alike, the cell-type-specific impact of *SETD1A*^+/-^ LoF has been supported by two mouse studies ^17,19^. Therefore, it is important to study *SETD1A* function in different models and different cell types. Since many mutations identified in *SETD1A* are associated with SCZ and NDDs, it is also important to study the mutation-specific impact of *SETD1A* related to different diseases.

Despite methodological differences, the consistent observation across studies is that *SETD1A*^+/-^ LoF mutations significantly impact dendritic architecture and synaptic function. This convergence suggests that SETD1A plays a crucial role in maintaining neuronal structural integrity and synaptic function. Future multi-modality functional assay of neurons carrying *SETD1A* LoF mutations especially in the context of other neurodevelopmental disorder risk genes on a large scale will further establish the functional role of *SETD1A* in contributing neuropsychiatric and neurodevelopmental disorders ^59^. Given the association of various *SETD1A* mutations with different NDDs and SCZ, it is evident that understanding the biological functions of SETD1A in different neural contexts is essential for unraveling the pathophysiology of these conditions. Our study highlights the crucial role of SETD1A in neuronal development and synaptic function, with implications for understanding the pathophysiology of SCZ and NDDs. Further research is needed to elucidate the precise mechanisms and develop targeted therapies.

## Acknowledgment

The Child Health Institute of New Jersey is supported in part by the Robert Wood Johnson Foundation (RWJF grant #74260). We also thank other members of the Pang laboratory and collaborators of the *SETD1A* project for their support and valuable comments.

## Author contributions

X.S. conducted the main experiments, including cell culture, electrophysiology, morphological and genomics analysis. H.Z., Q.Y. and Y.H. conducted the CRISPR/Cas9 gene targeting, and related analyses and provided conceptual input on experimental design. L.W., T.L., L.C., E.L., S.Z., and K.M. conducted part of the analysis. J.L. and A.K. conducted eSNP-Karyotyping analysis. H.S., G.M., J.D. and Z.P.P conceived the project. X.S., J.D., and Z.P.P. wrote the paper with input from all authors.

## Data and code availability

The scripts for running the RNA-Seq processing and eSNP-karyotyping pipeline are provided at https://zenodo.org/records/13840811. Raw and processed RNA-Seq data are available at GEO XXX.

